# A quantitative PCR assay for detection of the mycotoxigenic plant pathogen and food spoiling mold *Paecilomyces niveus* in fruit, food, and soil

**DOI:** 10.1101/2023.02.18.529073

**Authors:** Tristan W. Wang, David A. Strickland, Yasmin Haredy, Kerik D. Cox, Kathie T. Hodge

## Abstract

The postharvest fruit pathogen *Paecilomyces niveus* produces ascospores that can survive some pasteurization temperatures, spoil fruit products, and contaminate them with patulin, an FDA-regulated mycotoxin. Preventing *P. niveus* from entering food systems requires a robust detection method to effectively determine sources of *P. niveus* spoilage and disease inoculum. We designed a new robust and culture-independent detection method using species-specific primers (PnPATf/r) based on the patK gene, encoding a 6-methylsalicylic acid synthase, in *P. niveus*, for use in a rapid qPCR assay. Primer specificity was validated using 24 different *P. niveus* isolates and 16 other important food spoilage and fruit pathogenic fungi. The threshold for detection of qPCR was 18 genome equivalents. To further validate our new detection method, we demonstrate its use in detecting *P. niveus* in infected fruits, infested soils and ciders, and in fruit arising from apple blossoms sprayed with a *P. niveus* spore suspension. Results from this study may help fruit producers address spoilage and patulin contamination by this food spoiling fungus.

**Highlights:**

- New primers specific to *Paecilomyces niveus* (PnPATf/r) were developed based on the patK gene
- A qPCR assay to detect *P. niveus* was validated, and shown to be able to detect quantities of *P. niveus* DNA as low as 18 genome equivalents
- The new qPCR assay was used to investigate the ability of *P. niveus* ascospores to infect strawberry fruits and enter apple fruits through apple blossom infestation

## 1. Introduction

*Paecilomyces* species are among the most widely encountered heat resistant molds (HRM) in processed fruit products (Snyder et al., 2019). *Paecilomyces niveus* Stolk & Samson (*Byssochlamys nivea* Westling) in particular can not only produce ascospores that can survive lower temperatures of pasteurization, but can also contaminate food products with patulin, an FDA-regulated mycotoxin (Frisvad 2018; Affairs 2020). This ascomycete fungus can also infect and grow in many fruits, causing the postharvest disease Paecilomyces rot; processing fruits infected with Paecilomyces rot can lead to product infested with *P. niveus* ascospores and juice contaminated with patulin (Biango-Daniels et al., 2019; Biango-Daniels & Hodge, 2018). Spores that persist through processing can germinate and continue to produce a variety of mycotoxins including byssochlamic acid, byssochlamysol, and mycophenolic acid (Mori et al., 2003, 2003; Puel et al., 2005).

*Paecilomyces niveus* is an extremotolerant fungus that can grow in environments with low oxygen, high acid levels, and low water activity (Affairs, 2020; Silva & Evelyn, 2020). Previous work has explored strategies of reducing *P. niveus* spoilage inoculum in fruit products, employing methods including ultraviolet light, hydrostatic pressure, and thermal treatment (Ferreira et al., 2009; Menezes et al., 2020; Palou et al., 1998). However, intense processing and treatment may degrade food quality and nutrition (Van Boekel, 2008). Detecting and preventing *P. niveus* from entering food systems presents a viable strategy for reducing spoilage and patulin contamination by this fungus while also preserving food quality.

It has been assumed that *P. niveus* spoilage inoculum originates from environmental sources and previous surveys suggest that the fungus is a common soil microbe (Biango-Daniels & Hodge, 2018; Put, 1964; Tournas, 1994). The fungus has been found in various fruit products including apple juices and in naturally infected fruits (Khokhar et al., 2019; Santos et al., 2018). Traditional culturing techniques and morphological identification are laborious and time-consuming and require taxonomic expertise. In addition, the sporadic nature of food spoilage presents a challenge for spoilage prevention strategies that rely on growing out food spoiling agents. We sought to develop a robust detection system to identify and quantify *P. niveus* DNA in industrially relevant environments.

We developed a highly specific primer pair based on the single copy patK gene, encoding a 6-methylsalicylic acid synthase, to detect *P. niveus* in fruits, soils, and foods (Banani et al., 2016). We then validated and applied our qPCR-based detection method to two industrially relevant situations: 1) infected strawberries, and 2) fruits that developed from infested apple blossoms. A reliable detection method for *P. niveus* can contribute to strategies for reducing spoilage and contamination in downstream food processing.

## 2. Materials and methods

### 2.1 Specific primers were based on the patulin polyketide synthase gene

Low-coverage genomes of 23 isolates of *P. niveus*, sampled from across New York State, were aligned to the genome of *P. niveus* isolate CO7 for visualization of polymorphic sites within the patK gene, a component of the patulin secondary metabolite gene cluster (Biango-Daniels et al., 2018). Primers PnPATf and PnPATr (Table 1) were selected from appropriate regions using PrimerQuest software from Integrated DNA Technologies. Primer specificity was validated in silico using Primer-BLAST searches on related fungal species.

**Table 1.**
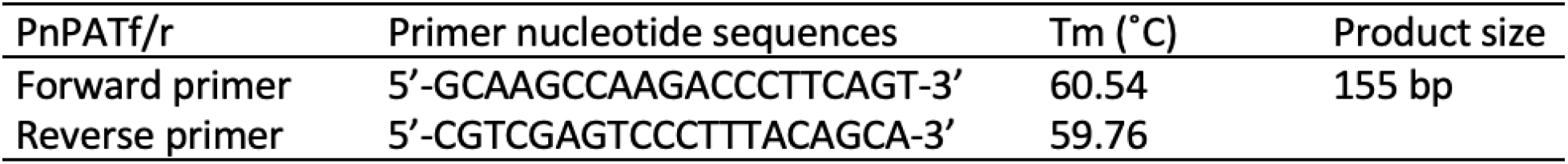
Primers developed in this study for qPCR assays

### 2.2 Culture preparation and DNA extraction

Fungal isolates used to evaluate primer specificity (Tables 1 and 2) were obtained from laboratory culture collections of the Hodge Lab at Cornell University and the USDA-Agricultural Research Service (NRRL) culture collection. Fungi were cultured on sterile cellophane overlaid on Potato Dextrose Agar (PDA; Criterion) at 25°C for 2 weeks in the dark, with the exception of *Aspergillus flavus* AF-36, which was grown at 33°C. Genomic DNA was isolated using the DNeasy PowerSoil Pro Kit (Qiagen) following the manufacturer’s instructions.

**Table 2.** 16 fungal species and 24 *Paecilomyces niveus* isolates used in validation of primers PnPATf and PnPATr, and presence or absence of PCR product. * = Food spoiling agent, ** = fruit pathogen

| Species | Strain | Source of isolation | PCR product |
| --- | --- | --- | --- |
| *Aspergillus flavus* * | AF-36 | Cotton fields, AZ | - |
| *Aspergillus terreus* * | NRRL 269 | Food, fruit (dates), CA | - |
| *Botrytis obtusa* ** | 97017 | Unknown substrate | - |
| *Mucor piriformis* ** | NRRL 3637 | Soil, Austria | - |
| *Neosartorya fisherii* * | 84-142 | Unknown substrate | - |
| *Neosartorya glabra* * | NRRL 3434 | Soil, Japan | - |
| *Neosartorya lanciniosa* * | NRRL 4076 | Hay, England | - |
| *Neosartorya spinosa* * | NRRL 32569 | Soil, Brazil | - |
| *Talaromyces bacillisporus* * | NRRL 1025 | Begonia leaf, NY | - |
| *Talaromyces islandicus* * | NRRL 1036 | Unknown substrate, South Africa | - |
| *Paecilomyces fulvus* * | 7 | Spoiled food | - |
| *Paecilomyces niveus* | 109-21 | Orchard soil, NY | + |
|  | 107-1 | Orchard soil, NY | + |
|  | MC4 | Residential soil, NY | + |
|  | CO7 | Culled apples, NY | + |
|  | 102-1 | Research station soil, NY | + |
|  | 100-10 | Orchard soil, NY | + |
|  | CF6 | Corn field soil, NY | + |
|  | 108-11 | Orchard soil, NY | + |
|  | CFO13 | Corn field soil, NY | + |
|  | CC8 | Compost, NY | + |
|  | 111-1 | Orchard soil, NY | + |
|  | 112-22 | Soil, NY | + |
|  | BY11 | Barn yard Soil, NY | + |
|  | 146-311 | Residential soil, NY | + |
|  | EP1 | Rhizosphere soil, unknown location | + |
|  | 141-11 | Farm soil, NY | + |
|  | 104-22 | Orchard soil, NY | + |
|  | 110-3 | Orchard soil, NY | + |
|  | 125-31 | Farm market soil, NY | + |
|  | AF01 | Alfalfa field soil, NY | + |
|  | 101-3 | Research farm soil, NY | + |
|  | 106-3 | Orchard soil, NY | + |
|  | KRA4 | Food facility, unknown location | + |
|  | 140-11 | Farm soil, NY | + |
| *Paecilomyces variotii* * | 103-2 | Soil, NY | - |
| *Penicillium expansum* ** | 94222 | Unknown substrate | - |
| *Penicillium griseofulvum* ** | NRRL 2159A | Unknown substrate | - |
| *Penicillium carneum* ** | NRRL 25170 | Unknown substrate | - |
| *Penicillium italicum* ** | KH1 | Diseased clementine, NY | - |

### 2.3 Ascospore extraction and soil and cider infestation

Spores for the qPCR assay were harvested from sterile cellophane overlaid on PDA. After four weeks, the surface of the colony was covered with 2mL of autoclaved deionized water and agitated to release asci and ascospores. To remove hyphal fragments, the spore suspension was filtered through eight layers of cheesecloth and Whatman No. 1 filter paper. A haemocytometer was used to quantify spore concentration by counting both free ascospores and 8-spored asci. Spore suspension underwent centrifugation to increase spore concentration if needed.

Soil samples for the qPCR assay were collected from an apple orchard near Ithaca, New York, and apple cider was purchased from a local supermarket. 10^7^ ascospores in 10 µl of water was used to infest 100 mg samples of orchard soil and 100 µl aliquots of apple cider. Negative controls were treated similarly but with 10 µl sterile water. Our DNA extraction method was modified for *P. niveus* ascospore lysis: Microcentrifuge tubes containing soil or cider samples infested with *P. niveus* ascospores underwent a 2 hour heat treatment at 70°C in a water bath. Heat-treatment was immediately followed by alternating periods of intense bead-beating using a mini-beadbeater (Biospec Products, Bartlesville, OK) (5 × 60 second intervals at 50 RPMx100) and 2 minutes of rest to prevent overheating prior to DNA extraction with the DNeasy PowerSoil Pro Kit.

### 2.4 Fruit infection assays and *Paecilomyces niveus* quantification

For qPCR detection of *P. niveus* in infected fruits, Fuji apples and California Navel oranges purchased from a local supermarket were sanitized and then infected using infested toothpicks according to the method outlined by Biango-Daniels and Hodge (2018). Albion strawberry fruits were obtained from Cornell Orchards and were sanitized similarly. The strawberry fruits were wounded with a pipette tip about 1 cm deep and 10^5^ ascospores in water were pipetted into the wounds. All fruits were kept in moist chambers in the dark at 25°C. Roughly 400 mg of infected fruit flesh was extracted from the resulting lesions of infected fruits, 2 cm away from the point of inoculation. Diseased flesh was extracted from apple and orange fruits two weeks post-inoculation and from strawberry fruits one week post-inoculation. Fruit flesh was extracted directly from apple and strawberry lesions while the peel was removed before fruit flesh extraction of orange fruits. DNA was quantified using qPCR for DNA extracts from three Fuji apples, three California Navel oranges, and four Albion strawberry fruits. This process was repeated for control fruit samples that had been mock-inoculated with sterile toothpicks under the same sanitization, incubation, and extraction protocol (Biango-Daniels & Hodge, 2018).

Because this is the first study to investigate strawberry susceptibility to Paecilomyces rot, disease-causing ability was evaluated based on satisfaction of Koch’s postulates.

### 2.5 Apple blossom inoculations

To investigate the possibility that blossom infection may be one route by which *Paecilomyces niveus* invades developing apples, *P. niveus* ascospore suspension was sprayed onto apple blossoms at various phenological stages: 10% bloom, 100% bloom, and petal fall. Developing fruits were later tested for presence or absence of *P. niveus* using our qPCR assay.

A field trial was conducted at a Cornell AgriTech research orchard in Geneva, NY in 2020 and 2021. The experiment was performed in a high-density orchard of dwarf apple trees,’NY109’ (‘Firecracker’) on G.935 rootstocks, established in 2016. Trees were planted at approximately 2 meter in-row spacing and trellised to three wires. The planting received no pesticides throughout the trial years.

Treatments were made on 8 single-tree replications arranged in a randomized complete block design. Treatments were made to runoff with a 4-gallon HDPE backpack sprayer (Solo Inc., Newport News, VA), with application focus on the blossoms of each tree. Applications consisted of inoculum suspended in water amended with 0.02% Tween-80, with a *P. niveus* spore concentration of 1 × 10^5^ asci and ascospores per mL. Prior to application, inoculum was produced on PDA as described in Section 2.2. In 2020, ascospore suspensions were applied on 20 May (10% bloom), 27 May (100% bloom), and 2 June (petal fall). On September 28, 2020, half the fruits were harvested and stored at room temperature to allow for rot development, and the other half was frozen for later evaluation. In 2021, ascospore suspensions were applied on 6 May (10% bloom), 10 May (100% bloom), and 17 May (petal fall). On September 14, 2021, apple fruits were harvested and stored under the same conditions as in the previous year.

Three batches of five apples were blended for each set of apples: 10% bloom, 100% bloom, petal fall, and uninoculated apples. This process was repeated for both the 2020 and the 2021 growing season. 400 mg of apple puree of each homogenized batch underwent DNA extraction for a total of 24 samples.

### 2.6 PCR and qPCR amplification conditions

End-point PCR reaction mixture contained 5 µl of 5X Q5 reaction buffer, 0.5 µl 10 mM dNTPs, 1.25 µl 10 µM of each primer, 1.25 µl template DNA, 0.25 µl of Q5 High-Fidelity DNA polymerase (New England Biolabs, Ipswich, MA, USA), and 15.5 µl nuclease-free water. PCR amplification was carried out on a PTC-200 Peltier Thermal Cycler (BioRad, Toronto, Ontario). The initial denaturation occurred at 98°C for 30s, followed by 26 cycles of denaturation at 98°C for 10 s, annealing at 64°C for 30 s, and extension at 72°C for 45 s. A final extension at 72°C occurred for 2 minutes. ITS4 and ITS5 Primers amplifying the internal transcribed spacer were used as a positive control (White et al. 1990). PCR products were then electrophoresed with TBE in 1.0% agarose gels (7V/cm, 60min) stained with Gel-Red (Biotium Inc., Hayward, USA).

*Paecilomyces niveus* DNA from extracted fruit, soil, and food was quantified using the CFX Connect Real-Time PCR Detection system (Bio-Rad, Hercules, CA, USA). qPCR reaction mixtures contained 25 µl 2x Power SYBR Green PCR Master Mix (Applied Biosystems, Madrid, Spain), 1 µl 10 µM PnPATf primer, 1 µl 10 µM PnPATr primer, 10 µl of eluted fungal DNA, and 13 µl deionized water (50 µl total). qPCR was performed on the CFX Connect Real-Time PCR System (Bio-Rad, Hercules, CA, USA) programmed to hold at 95°C for 10 minutes and then to complete 40 cycles of 95°C for 15 s (denaturation), 60°C for 30 s (annealing), and 72°C for 45 s (extension). Reactions were done in triplicate and results were measured by quantification cycle (Cq).

### 2.7 Standard curve

A ten-fold serial dilution of *P. niveus* CO7 DNA ranging from 7 to 7 × 10^−4^ ng/µl was generated and quantified using a Nanodrop-8000 Spectrophotometer (Thermo Scientific, Wilmington DE, USA) (260/280 ratio of 1.89). These DNA concentrations were used to generate a standard curve by qPCR by plotting genome equivalents against Cq values.

## 3. Results

### 3.1 Testing the PnPATf and PnPATr primer pair for species specificity

End-point PCR was used to validate primer specificity in vitro for a set of 19 fungal species and 24 isolates of *Paecilomyces niveus* (Table 2). These fungal isolates include apple pathogens (*Botrytis obtusa, Mucor piriformis, Penicillium* spp., *Venturia inaequalis*) and a citrus pathogen (*Penicillium italicum*). Primers were also tested on a variety of food spoilage fungi (*Aspergillus* spp., *Neosartorya* spp., *Talaromyces* spp.) and three closely related species (*Thermoascus crustaceou*s, *Paecilomyces fulvus, Paecilomyces variotii*). The 155 bp PCR product amplified by primers PnPATf and PnPATr was observed only in reactions including any of the 24 *P. niveus* isolates, as confirmed by gel electrophoresis.

### 3.2 Determining sensitivity and amplification efficiency of SYBR green qPCR assay

The detection limit of the qPCR method was established by testing a dilution series of *Paecilomyces niveus* isolate CO7 genomic DNA. Concentrations ranging from 7 ng/µl to 0.7 pg/µl were tested in triplicate. The qPCR assay was able to detect *P. niveus* DNA when there were only 7 pg of *P. niveus* DNA (about 18 genome equivalents) in the 50 µl reaction (Cq value of 35.04 ± .9344) (Figure 1). The amplification efficiency of the qPCR method was 87.28% based on the equation ε = 100 × (10^−1/slope^ -1) (Broeders et al., 2014). There was no detection in any of the negative controls.

**Figure 1.**
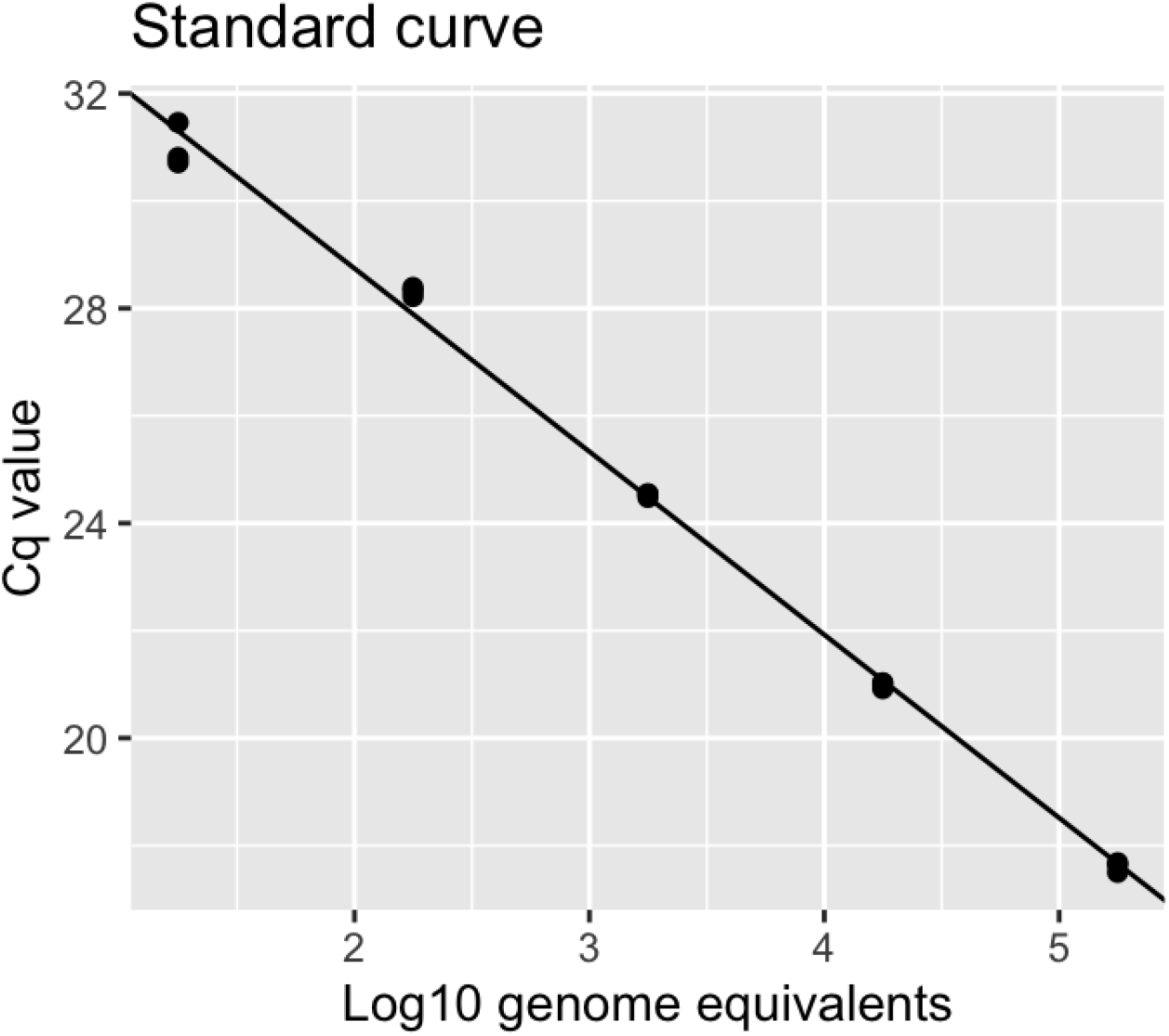
qPCR amplification of *Paecilomyces niveus* DNA (concentrations ranging from 7 ng/µl to 0.7 pg/µl) using log10 values of *P. niveus* genome equivalents plotted against quantification cycle (Cq). Standard equation: Cq = -3.4093x + 35.56.06, R^2^ = .9965 (efficiency = 96.48%)

### 3.3 Quantification of Paecilomyces niveus DNA in fruit, soil, and cider

The qPCR assay was applied to eluted DNA extracted from developing lesions of fruits infected with *P. niveus*. DNA was extracted from 3 infected apple fruits, 3 infected orange fruits, and 4 infected strawberry fruits. Control fruits were treated similarly but without the presence of *P. niveus. Paecilomyces niveus* DNA was detected in all infected apple and orange fruits and in three of the four strawberry fruits (Table 3). No results were detected in the control fruits. The SYBR green assay was then used to quantify *P. niveus* CO7 DNA from 3 soil and apple cider samples infested with *P. niveus* ascospores. Cq values of DNA extract obtained from puree of apple fruits that developed from infested blossoms were negative for the 10% bloom, 100% bloom, and petal fall trials.

**Table 3.** Average Cq values from qPCR amplification of *Paecilomyces niveus* CO7 DNA from apple and orange fruits, food, and soil (n = 3) and from strawberries (n = 4). *P. niveus* DNA was not detected in one sample from an infected strawberry. DNA was extracted from fruit flesh of infected apple and orange fruits two weeks post-inoculation, from ciders and soils infested with *P. niveus* hyphae, and cider and soils infested with 10^7^ *P. niveus* ascospores.

| Substrate | *Paecilomyces niveus* treatment | Cq value (average ± sd) |
| --- | --- | --- |
| Infected apple fruit | Hyphae | 20.80 ± 1.27 |
| Infected orange fruit | Hyphae | 17.78 ± 2.26 |
| Infected strawberry fruit | 10<sup>5</sup> ascospores | 22.49 ± 10.37 |
| Infested cider | 10<sup>7</sup> ascospores | 19.41 ± 1.44 |
| Infested soil | 10<sup>7</sup> ascospores | 27.34 ± .13 |

## 4. Discussion

This study presents a new qPCR approach to detect *Paecilomyces niveus* spores or mycelium in fruit, soil, and fruit matrices. We designed a pair of primers (PnPATf and PnPATr) specific to a 155-bp locus within the patK gene. We also developed a straightforward DNA extraction protocol to break tough-to-lyse *P. niveus* ascospores to improve detection at lower levels. The primer pair was validated in silico and in vitro through end-point PCR on a sample of 24 *P. niveus* isolates and 16 food spoilage fungi, fruit pathogens, and close relatives. PCR products were obtained only from the 24 *P. niveus* isolates. From qPCR amplification of pure *P. niveus* CO7 DNA, a standard curve (R^2^ = .99) was established,showing this method was able to detect and quantify *P. niveus* DNA across a range of conditions including low concentrations of DNA, on par with previous qPCR work on food spoiling fungi (Gil-Serna et al., 2009; Panek & Frą c, 2018). Importantly, our newly developed primers did not detect the close relatives *P. fulvus* and *P. variotii*, nor various patulin-producing members of the Eurotiales including the *Penicillium expansum, P. carneum*, and *P. griseofulvum*. This is in part due to an absent or divergent patK (Puel et al., 2007; Urquhart et al., 2018). Compared to a growing array of methods for detection of food spoilage fungi including the use of biosensors, metabolic profiles, phenotypic assays, and other molecular techniques like loop-mediated isothermal amplification, our method is promising in that it is both robust and accessible in not relying on highly sophisticated equipment (Panek & Frą c, 2019; Pertile et al., 2020; Santana Oliveira et al., 2019). Future research may focus on detecting multiple thermotolerant food spoiling or patulin-producing fungi via a multiplexing approach, for use in industrially and agriculturally relevant environments.

We used the primer pair PnPATf/PnPATr in our newly developed qPCR assay for detection of *P. niveus* DNA in infected fruits, and in infested soil and cider. Higher Cq values from cider and soil samples infested with 1×10^7^ ascospores, compared to Cq values from pure *P. niveus* DNA extract suggest some level of inhibition from the food and soil matrices during DNA extraction and PCR. We found that the tough spore walls, coupled with their small size (ascus diameter < 10 µm), presented a challenge for lysis. Various spore pre-treatments were evaluated to optimize DNA yield, including use of NaOH, phenol-chloroform, and cell wall digesting enzymes. Using our bead-beating and heat treatment protocol, we were able to detect *P. niveus* in all samples of infested soil and cider.

*Paecilomyces niveus* has several potential routes into food: through environmental sources like air and water, and by infection of fruits with Paecilomyces rot (Biango-Daniels et al., 2019). In this study we deployed our qPCR detection method to evaluate whether Paecilomyces rot can develop in apples after ascospore infestation of apple blossoms, and after ascospore infection of strawberry fruits. The fungus has been shown to be a wound-infecting pathogen of several rosaceous fruits, and to colonize the apple core during infection (Biango-Daniels & Hodge, 2018; Wang & Hodge, 2022). In several other postharvest fruit pathogen systems like moldy-core caused by *Alternaria alternata* and gray mold of strawberries caused by *Botrytis cinerea*, disease inoculum is presumed to enter fruits before fruit maturation and cause latent infections (Bristow et al., 1986; Reuveni et al., 2002). A previous report of natural Paecilomyces rot infections showed diffuse brown lesions suggesting the presence of internal rot (Khokhar et al., 2019). Contrary to our hypothesis that Paecilomyces rot can result from blossom infestation, our qPCR results did not detect significant *P. niveus* DNA from apples that developed from a small sample of previously infested blossoms. We speculate that because Paecilomyces rot development is slower than some other post-harvest diseases like Blue mold in apples, not enough *P. niveus* biomass accumulated for possible detection. It is also possible that low disease incidence contributed to the lack of detection. We did observe lesion development in strawberry fruits after *P. niveus* ascospore inoculation, thus this the first study to demonstrate the susceptibility of strawberries to Paecilomyces rot.

Our qPCR assay for detection and quantification of *P. niveus* DNA allowed us to ask questions and test hypotheses regarding *Paecilomyces niveus* disease biology. This study provides a tool to help determine sources of *P. niveus* spoilage and disease inoculum, an important first step to combating spoilage and patulin contamination by this notorious fungus. Incorporating results from this study can help aid the development of comprehensive preventative strategies for spoilage and mycotoxin contamination in foods.

## Acknowledgements

We would like to thank the following people: Mykala Robertson and Marvin Pritts for providing fruits for inoculation, Gillian Turgeon and Jennifer Gonzalez for their expertise in genomics and protoplast generation, Jo Ann Asselin, Mary Ann Karp, Wei Zhang, Melanie Filiatraut, and Juliana Gonzalez Tobon for their aid in our qPCR applications, and Maryanne McCloskey for contributing to DNA extraction.

